## Supplementary material for "GPCR binding and JNK3 activation by arrestin-3 have different structural requirements": S1

**Supplementary Information**

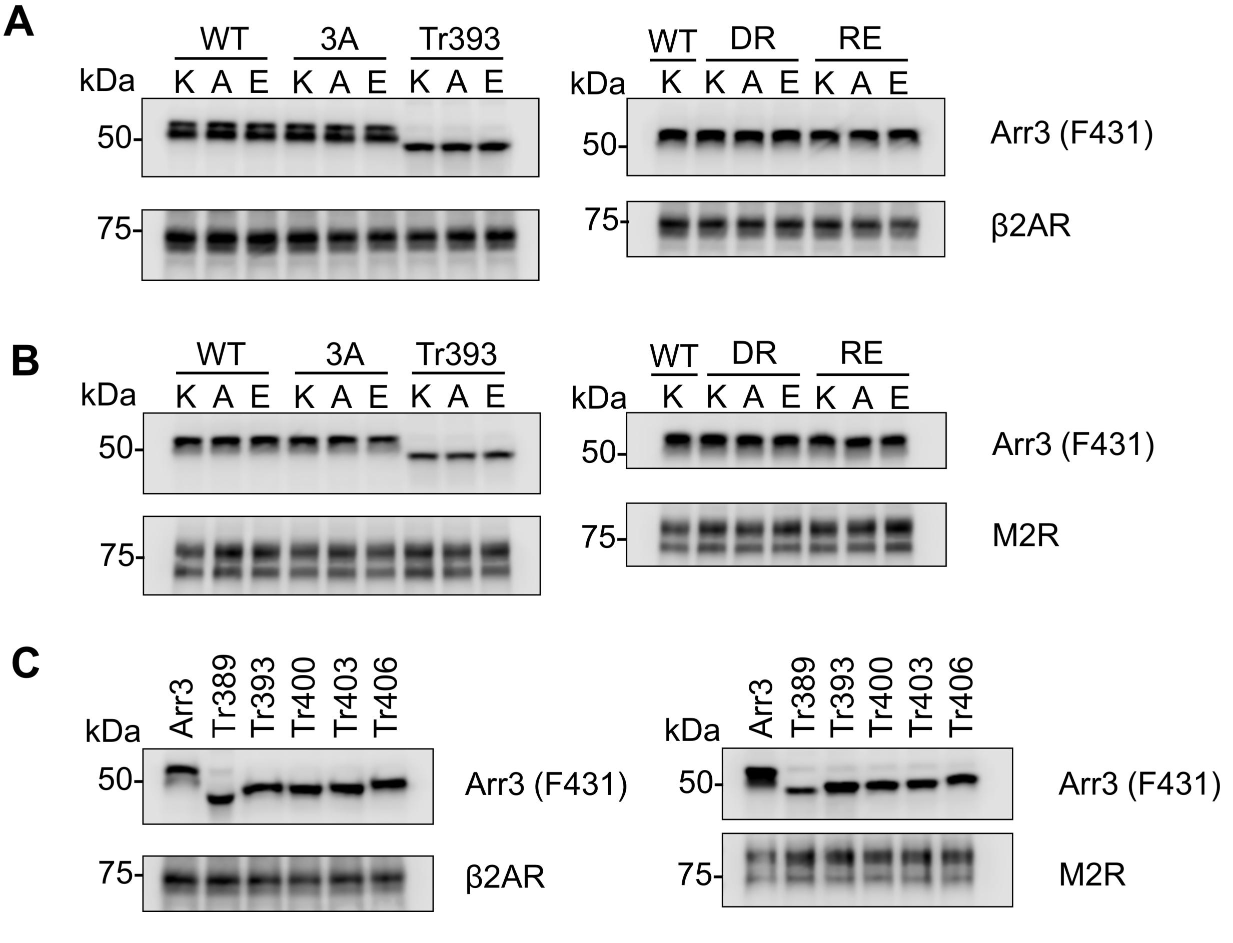


**Fig. S1**. **Expression of SmBiT-arrestin-3 and LgBiT-GPCRs used in nanoBiT assay.** Arrestin-3 and receptors were detected by western blot with anti-arrestin F431 [1] and anti-HA (#3724, Cell Signaling Technology) antibodies, respectively. **A.** Samples shown in Fig. S3. **B.** Samples shown in Fig. S4. **C.** Samples shown in Fig. 3.

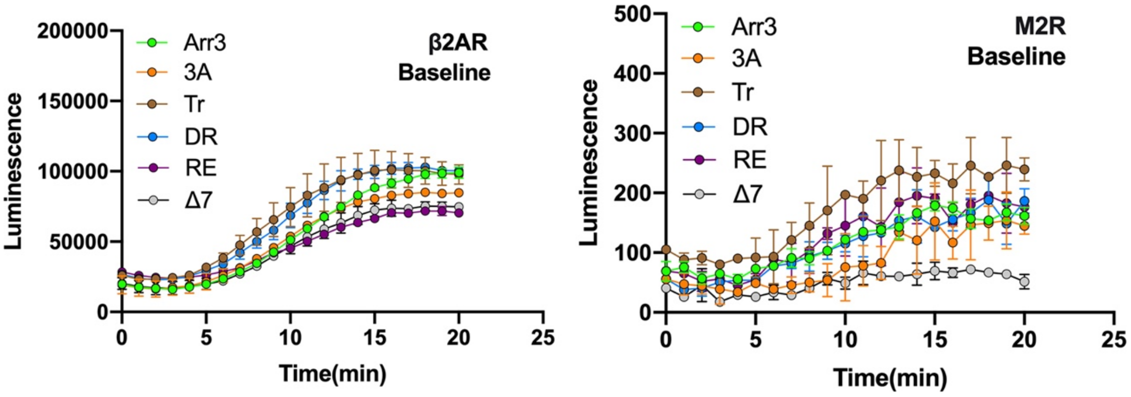


**Fig. S2**. **Basal luminescence (before agonist stimulation) in cells expressing LgBiT-tagged β2AR and M2R with SmBiT-tagged WT arrestin-3 and indicated mutants shown in Fig. 2.** Note that the signal with β2AR is much larger than with M2R. The most likely explanation is that β2AR has phosphorylation sites in the C-terminus, where LgBiT is fused, which binds in the cavity of the N-domain [2], close to the SmBiT fused to the N-terminus of arrestin-3. In contrast, M2R has phosphorylation sites necessary for arrestin binding in the third cytoplasmic loop [3,4], while the LgBiT is fused to its short C-terminus that does not have any phosphorylation sites.

A

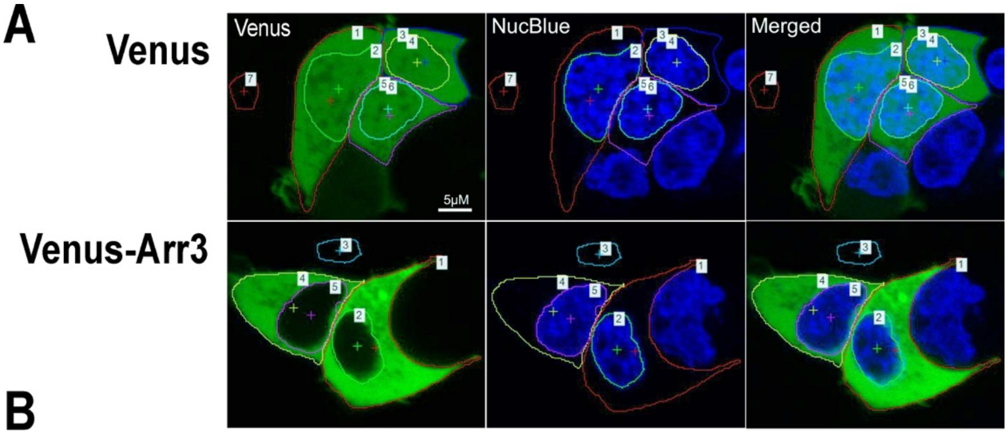


B

| Venus | | | | Green Fluorescence (488 nm) | | NET Fluorescence | | Normalization | |
|---|---|---|---|---|---|---|---|---|---|
| | | | | MEAN Intensity per pixel | Total Intensity | MEAN Intensity per pixel | Total Intensity | MEAN Nuclear Intensity per px/MEAN Total/px | Nuclear content, % of Total) |
| Cell# | Area# | Field | ROI, pixels | | | | | | |
| 1 | 1 | Total | 7506 | 75.04 | 563213 | 73.56 | 552104 | 1.0150 | 49% |
| | 2 | Nuclear | 3592 | 76.15 | 273535 | 74.67 | 268219 | | |
| 2 | 3 | Total | 3772 | 83.52 | 315030 | 82.04 | 309447 | 0.9950 | 51% |
| | 4 | Nuclear | 1935 | 83.07 | 160734 | 81.59 | 157870 | | |
| 3 | 5 | Total | 3797 | 77.29 | 293461 | 75.81 | 287841 | 1.0450 | 57% |
| | 6 | Nuclear | 2080 | 80.72 | 167904 | 79.24 | 164826 | | |
| Background | 7 | Background | 627 | 1.48 | 930 | - | - | | |

| Venus-Arr3 | | | | Green(GFP) | | NET Fluorescence | | Normalization | |
|---|---|---|---|---|---|---|---|---|---|
| | | | | MEAN Intensity per pixel | Total Intensity | MEAN Intensity per pixel | Total Intensity | MEAN Nuclear Intensity per px/MEAN Total/px | Nuclear content, % of Total |
| Cell# | Area# | Field | ROI | | | | | | |
| 1 | 1 | Total | 5229 | 125.31 | 655270 | 123.83 | 647531 | 0.3520 | 10% |
| | 2 | Nuclear | 1531 | 45.07 | 69005 | 43.59 | 66739 | | |
| 2 | 4 | Total | 3980 | 67.49 | 268630 | 66.01 | 262740 | 0.3730 | 18% |
| | 5 | Nuclear | 1890 | 26.12 | 49365 | 24.64 | 46568 | | |
| Background | 3 | Background | 512 | 1.01 | 518 | - | - | | |

**Fig. S3. Quantification of the subcellular distribution of arrestin-3 wild type and mutants.** HEK293 arrestin-2/3 KO cells were co-transfected with HA-ASK1, HA-JNK3α2 and either control (Venus) or indicated N-terminally Venus-tagged forms of arrestin-3 (to mimic the condition of the JNK activation). The images were collected from live cells 48 h post-transfection on the Olympus confocal microscope, as described in Methods. The images were analyzed for the intensity of the green (488 nm) fluorescence using NIS-elements software. **(A)** Representative cells expressing Venus or Venus-tagged WT arrestin-3 (Venus-Arr3). The total cell area and the nuclear area (stained by NucBlue) are outlined representing Regions of Interest (ROI) marked by numbers for the analysis. A cell-free area is used to determine the background (#7 in Venus and #3 in Venus-Arr3). **(B)** The software provides measurements of ROI in pixels, fluorescence intensity per pixel and sum of intensity for the entire ROI. NET values are measurements after the subtraction of the background. To estimate a relative enrichment of the arrestin-3 protein in the nucleus, we used the mean nuclear fluorescence intensity per pixel normalized to the mean total intensity per pixel (to account for the differences in the levels of transfection among the cells). To estimate the overall distribution of the arrestin-3 proteins between the nucleus and the cytosol, we calculated the percentage of the total expressed protein residing in the nucleus (nuclear fluorescence as % of the total fluorescence).


1. Vishnivetskiy, S.A., Zhan, X., Chen, Q., Iverson, T.M., and Gurevich, V.V. (2014). Arrestin expression in E. coli and purification. Curr Protoc Pharmacol *67*, Unit 2.11.11-19.
2. Lee, Y., Warne, T., Nehmé, R., Pandey, S., Dwivedi-Agnihotri, H., Chaturvedi, M., Edwards, P.C., García-Nafría, J., Leslie, A.G.W., Shukla, A.K., and Tate, C.G. (2020). Molecular basis of β-arrestin coupling to formoterol-bound β(1)-adrenoceptor. Nature *583*, 862-866. 10.1038/s41586-020-2419-1.
3. Nakata, H., Kameyama, K., Haga, K., and Haga, T. (1994). Location of agonist-dependent-phosphorylation sites in the third intracellular loop of muscarinic acetylcholine receptors (m2 subtype). Eur J Biochem *220*, 29-36.
4. Lee, K.B., Ptasienski, J.A., Pals-Rylaarsdam, R., Gurevich, V.V., and Hosey, M.M. (2000). Arrestin binding to the M2 muscarinic acetylcholine receptor is precluded by an inhibitory element in the third intracellular loop of the receptor. J Biol Chem *275*, 9284-9289.
